# Transcript-Capture sequencing enriches mRNA of *Mycobacterium tuberculosis* from host samples

**DOI:** 10.1101/2025.10.28.685133

**Authors:** Eleanor I. Lamont, Richard M. Jones, Jessica Assadi, Robert Morrison, Taeksun Song, Xiang Yu, Danielle M. Weiner, Laura E. Via, Jill Winter, Shuyi Ma, Robert J. Wilkinson, Clifton E. Barry, David R. Sherman

## Abstract

Bacterial gene expression from sites of infection are poorly studied due to low levels of bacterial mRNA present in clinical samples. Here, we develop Transcript-Capture Seq, which uses customizable biotinylated probes generated in-house to enrich bacteria-specific RNA from host samples before Next Generation Sequencing (NGS). This method results in a >200-fold increase in bacterial mRNA reads from mixed samples and allows analysis of the complete bacterial transcriptome from clinical samples. We apply Transcript-Capture to models of tuberculosis (TB) infection as well as sputum samples from TB patients. TB patients exhibit unexplained heterogeneity in disease progression, and the activity of *Mycobacterium tuberculosis* (Mtb) has been proposed to affect treatment response. By applying Transcript-Capture to sputum samples collected from TB patients we generate the first complete *in vivo* bacterial transcriptome of Mtb via NGS. Mtb from patient sputa shows upregulation of genes involved in host lipid utilization and zinc limitation, as well as a similar gene expression profile to Mtb log phase growth *in vitro*. Applying Transcript-Capture to clinical sputa provides a snapshot of bacterial activity directly from human patients and can be used to investigate the physiological state of bacteria surviving *in vivo*.

**GRAPHICAL ABSTRACT:** 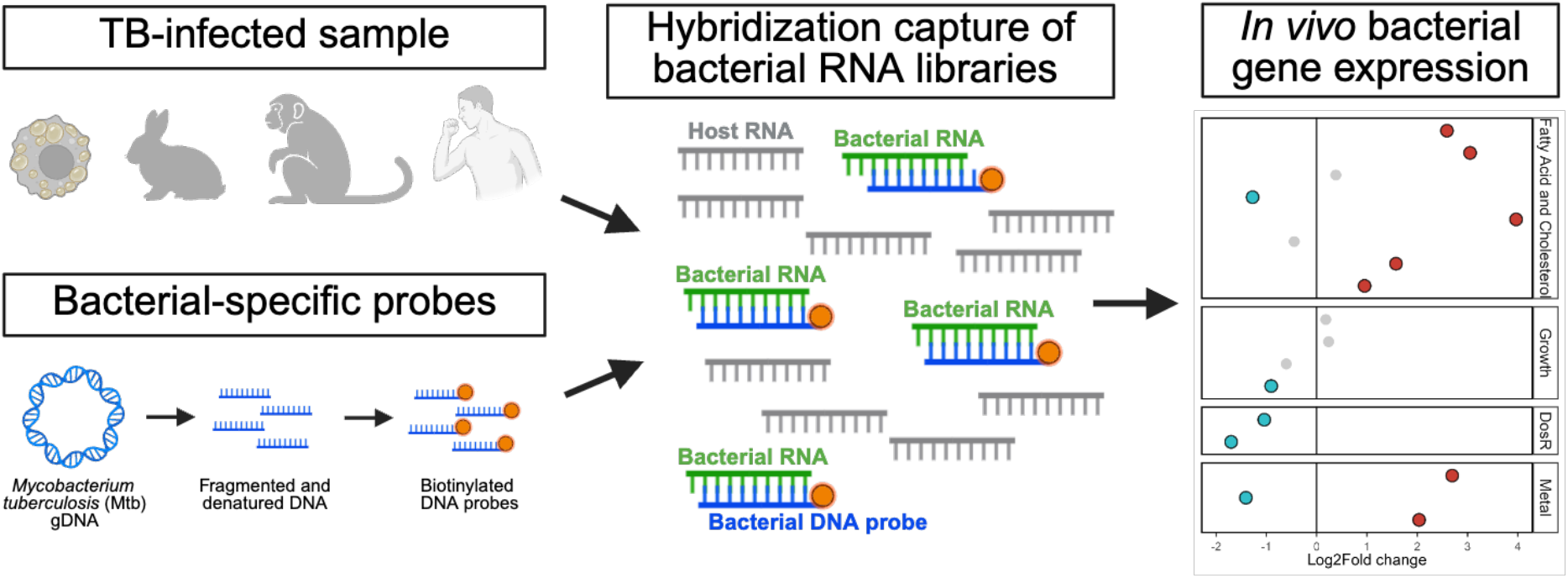

## INTRODUCTION

The mRNA of bacterial pathogens within host environments provides a unique perspective into bacterial physiological state that may help us understand and combat infectious diseases. However, bacterial activity *in vivo* is often poorly characterized due to technical challenges of processing the extremely low levels of bacterial material present in most samples [1–4]. Several methods have been developed to enrich small amounts of bacterial mRNA out of mixed samples for Next Generation Sequencing (NGS) applications, including using target amplification with commercially synthesized PCR probes that hybridize and extract matching sequences in an RNA sequencing library [5–9]. Although these probes have successfully amplified bacterial RNA from murine and eukaryotic infected cells, this strategy remains costly due to the need for custom probe design and synthesis, captures a limited number of genes, and has not yet been validated in clinical samples.

Genome Capture Sequencing (GenCap-Seq) is a method of nucleic acid enrichment that was recently developed to sequence bacterial DNA in sputum from people living with cystic fibrosis [10,11]. This targeted enrichment is performed using lab-generated biotinylated DNA probes that hybridize to and enrich for DNA of specific bacterial strains from metagenomic sequencing libraries. By making probes in-house directly from the bacterial genome of interest, this protocol is cheaper than commercial synthesis methods, produces longer probes, and covers the entire bacterial genome. GenCap-Seq has been successfully used to generate whole-genome sequences from multiple species out of clinical samples such as sputum, nasal, fecal, and skin swabs that are composed mostly of human DNA and often contaminated with off-target bacteria [10,11].

Building on the GenCap-Seq strategy of generating probes directly from the bacterial genome, we have developed Transcript-Capture Seq, which uses biotinylated probes to enrich bacterial-specific RNA from host samples. Our workflow allows RNA sequencing at sufficient depth to interrogate complete bacterial transcriptomes from samples containing low levels of bacterial RNA in a large background of host material.

We have developed this protocol for use with *Mycobacterium tuberculosis* (Mtb) RNA from Mtb-infected host samples. Tuberculosis (TB) is responsible for over 10 million cases of active disease each year and patients exhibit unexplained heterogeneity in disease progression and treatment response. Most patients are cured well before the end of the standard 6-month antibiotic regimen while ∼5% of patients need longer drug therapy to prevent treatment failure or relapse [12–15]. Bacterial activity likely plays a key but unexplored role in treatment efficiency and the physiological state of Mtb *in vivo*, which can be analyzed via gene expression, is of critical interest for improving TB therapy [13,16,17]. However, NGS approaches have had limited success in directly sequencing Mtb RNA from host samples due to an abundance of contaminating host and microbiome RNA [18]. Here, we apply Transcript-Capture to *in vitro* and animal models of TB infection. We further assess Mtb from sputum samples of TB patients and generate complete transcriptomes of Mtb expectorated from the human lung. Our results indicate that Transcript-Capture should be useful in many other circumstances in which signal-to-noise concerns limit the utility of transcriptomics.

## MATERIALS AND METHODS

### Sample preparation and collection

#### Spiked samples

Cells of the human monocytic cell line THP-1 (ATCC #TIB-202) were spiked with the Mtb strain H37Ra and used as spiked samples to optimize Transcript-Capture. THP-1 cells maintained in RPMI medium were pelleted and resuspended in TRIzol (Invitrogen) at a concentration of 1×10^6^ cells/mL. 1mL THP-1 aliquots were added to lysing matrix B tubes (MPbio). The Mtb strain H37Ra was grown to log phase (OD_600_ = 0.2) in Middlebrook 7H9 medium (Sigma) with 10% OADC (Sigma), 0.2% glycerol, and 0.05% Tween 80 at 37°C. The THP-1 aliquots were spiked with between 1×10^2^ and 1×10^6^ cells of H37Ra directly before RNA extraction.

#### Caseum mimic samples

Caseum mimic was generated from THP-1 cells as previously described [19]. THP-1 cells were cultured in RPMI 1640 supplemented with 10% FBS and 2 mM L-glutamine. Cells were seeded at a density of 1×10^6^ cells/mL on 150 mm dishes and differentiated with 100 nM phorbol 12-myristate 13-acetate overnight. Lipid body formation was induced through the addition of 0.1 mM stearic acid dissolved in ethanol. After 24 hr, foamy macrophages were detached using 5mM EDTA, washed with PBS, and collected via centrifugation. The pellets were frozen and thawed three times using dry ice, then incubated at 75°C to denature proteins. Surrogate caseum pellets were then rested at 37°C for three days to stabilize and stored at −20°C until use. To generate Mtb samples adapted to caseum mimic, cultures of the Mtb strain HN878 were grown to an OD_600_ of 0.6-0.9. The culture was centrifuged and resuspended in water to an OD_600_ of 0.8. This bacterial cell suspension was added to the caseum mimic pellets in a ratio of 2:1 (vol/wt). Suspensions were briefly homogenized with 1.4 mm zirconia beads. Mtb was left to adapt to the surrogate matrix for 7-28 days at 37°C. Colony-forming units (CFU) were determined at each timepoint using dilution plating onto 7H10 solid media. Final burdens of Mtb after adaptation remained near 1 x 10^8^ cells.

#### Animal samples

All live vertebrate procedures were performed in accordance with the recommendations of the Guide for the Care and Use of Laboratory Animals of the National Institutes of Health and approved by the NIAID Animal Care and Use Committee in Protocol LCIM-3E (experimental tuberculosis in New Zealand White rabbits) or LCIM-9E (experimental tuberculosis in common marmosets) (Permit issued to NIH as A-4149-01). A female NZW rabbit (Charles River Labs, Fredrick MD) was infected by aerosol exposure to HN878 as previously described [20]. The rabbit samples used here were collected 26 weeks post-infection from an apparently healthy animal with stable weight > 4 kg yet bearing 7 computed tomography (CT)-identified cavities 5-22 mm in diameter. Marmoset samples were collected from a male marmoset infected by aerosol exposure to the Mtb strain CDC1551 as previously described [21,22]. The marmoset formed 5 lesions observed by [^18^F]-fluorodeoxyglucose positron emission tomography (PET) and CT by 4 weeks post infection; euthanasia and sample collection were performed 6 weeks post infection due to excessive weight loss. During necropsy, animal tissue was immediately apportioned into weighed Precellys CK14 Lysing Kit tubes (Bertin technologies) and vigorously mixed with cold TRIzol 2 cycles of 30 sec, at 5,000 rpm in a Precellys minilys tissue homogenizer. The processed samples were frozen on dry ice and stored at −80°C until RNA isolation.

#### Clinical sputum samples

The prospective, multicenter randomized clinical trial “Using biomarkers to predict TB treatment duration (PredictTB; Clinicaltrials.gov: NCT02821832)” has been previously described [12,23]. Participants gave written consent to participate in the trial though one of 5 local clinics in the Western Cape, South Africa overseen by their regional or university-based local ethical committees. Samples used here were collected during a study visit before the start of antibiotic treatment from HIV-negative patients 18-75 years old with body weight 35-90 kg who had not been treated for active TB within the past 3 years. A portion of sputum was preserved in TRIzol within 10 min of aspiration. The freshly produced sputum (1 to 1.5 mL) was mixed with an equal volume of TRIzol in a homogenizer or in a syringe fitted with a large gauge needle and dispensed into 2-mL tubes and frozen on dry ice/ethanol or liquid nitrogen. The specimens were held on dry ice for transport and storage at −80°C until RNA extraction. Mtb lineage was determined by analyzing whole genome sequences of Mtb strains isolated from sputum using the TB-profiler tool (v5.0.1) [24].

### RNA Extraction

RNA was extracted as previously described [25,26]. Samples stored in TRIzol were transferred to lysing matrix B tubes and shaken in a Bead Mill Homogenizer (OMNI International) with three 30 sec pulses and 30 sec rests on ice. Samples were centrifuged at max speed (13,000 x g), the supernatant was added to a phase lock tube (Invitrogen) containing 300µL chloroform, then the tube was inverted for two minutes. Samples were centrifuged at max speed and supernatant was added to a tube containing 300µL isopropanol and 300µL high salt solution (0.8M Na Citrate, 1.2M NaCl) and inverted to mix. Samples were stored at 4°C overnight to precipitate the RNA. After precipitation, samples were centrifuged at 4°C max speed for 10 minutes then pellets were washed once with 1mL cold 75% EtOH then resuspended in 80µL RNase free H_2_O. An initial DNase treatment was performed with TURBO DNase (Invitrogen). Total RNA was then purified using the RNeasy Micro Kit (Qiagen) following the manufacturer’s protocol for RNA Cleanup with a second on-column DNase treatment. After cleanup, rRNA was depleted using Mtb rRNA-specific biotinylated probes as described in [27]. rRNA probes were annealed to 150-250ng total RNA then rRNA strands were bound to streptavidin magnetic beads (NEB) for 5 min at room temperature and 5 min at 50°C. After annealing, the supernatant was removed, mixed with GuHCl binding buffer, and cleaned with Ampure XP beads (Beckman Coulter). RNA concentration and quality were measured with Nanodrop.

### Transcript-Capture probe generation

DNA used to make capture probes was extracted from the Mtb lab strain H37Rv using a chemical extraction procedure [28] to maintain intact DNA. DNA was sheared to ∼150bp in a COVARIS M220 and cleaned with 3X Ampure XP beads (Beckman Coulter). Sheared DNA was dephosphorylated with rSAP (NEB), denatured via heating, then biotinylated with biotin-11-ddATP (Revvity). Unbound biotin was removed using the Monarch PCR & DNA Cleanup Kit (NEB) following the manufacturer’s protocol for ssDNA purification. Biotinylated ssDNA was bound to Dynabeads MyOne Streptavidin C1 (Thermofisher). The manufacturer’s protocol was followed for bead washes with minor modifications: the bead-bound DNA was only washed once with a low salt formulation of Binding & Wash buffer and once with 0.1X SSC to remove any non-biotinylated DNA. Biotinylated DNA was dissociated from beads with 95% formamide and purified using the Monarch PCR & DNA Cleanup Kit (NEB) following the manufacturer’s protocol for ssDNA purification. The final biotinylated ssDNA probes were visualized on a dot blot with Streptavidin-AP conjugate (Roche) to confirm biotinylation and quantified by High Sensitivity ssDNA Qubit (Thermofisher). Detailed methods for probe preparation are provided in the supplementary material.

### RNA library preparation and Transcript-Capture

NGS libraries were prepared using NEBNext Ultra II RNA Library Prep Kit for Illumina (NEB) with NEBNext Multiplex Oligos for Illumina (Dual Index Primers Set 1) (NEB). Final libraries were quantified with Kapa Library Quantification kit for Illumina (Roche). Hybridization capture was performed using the xGen hybridization capture of DNA library protocol (IDT) with modifications: 3 to 5 samples (500ng cDNA each) were pooled with 25-50ng of custom DNA probe used for each pool. Detailed methods for Transcript-Capture are provided in the Supplementary Material. Samples were sequenced at the Northwest Genomics Center at the University of Washington on either Illumina NextSeq or Element Biosciences AVITI platforms. Only samples with successful sequencing of >1 million total reads were analyzed. FASTQ files are available in the NCBI Sequence Read Archive (SRA) under BioProject PRJNA1345083.

### Data analysis

RNA sequences were aligned to the Mtb genome [29,30] using the DuffyTools and DuffyNGS custom processing pipeline available at https://github.com/robertdouglasmorrison that uses Bowtie2 utilities. Mtb rRNA reads were excluded from analysis of reads aligning to Mtb, and only protein-coding genes were examined for differential gene expression (DEG) analyses. DEG was performed using the DuffyTools MetaResults function, which uses a combined result from five DEG tools (round robin, RankProduct, SAMs, EdgeR, and DeSeq2). Differential gene set expression analyses were performed using the DuffyTools MetaGeneSets function, which uses a combined result from five differential pathway tools (DensityPlots, RadarPlots, QuSage, Enrichment, and GSEA). Use of multiple differential expression analyses tools minimizes tool bias and provides a conservative metric of differential expression. To compare Mtb transcriptomes between the current study and previously published datasets, Spearman’s correlations were performed following quantile normalization of ranked gene expression values as described in [31]. For all statistical analyses, FDR-adjusted p-values < 0.05 were considered statistically significant. Graphs were made in R (v4.4.3) with the packages ggplot2 (v3.5.1), tidyverse (v2.0.0), ggcorrplot (v0.1.4.1), ggrepel (v.0.9.6), ggpubr (v0.6.1), and cowplot (v1.1.3).

## RESULTS

### Transcript-Capture uses bacteria-specific probes to enrich for Mtb RNA out of spiked samples

Here, we use custom-made DNA probes to enrich and sequence complete bacterial transcriptomes from mixed samples in which the majority of RNA derives from the host. Mtb-specific probes were generated for Transcript-Capture starting with a broth culture of the Mtb lab strain H37Rv. DNA extracted from H37Rv was fragmented to ∼150bp, denatured, and biotinylated to produce single-stranded biotinylated probes that cover the entire genome of Mtb (**Figure 1A**). Probes were then hybridized to rRNA-depleted RNA sequencing libraries after cDNA generation. Material that did not bind to the capture probes was washed away and the remaining library, enriched with Mtb cDNA fragments, was sequenced (**Figure 1B**).

**Figure 1.**
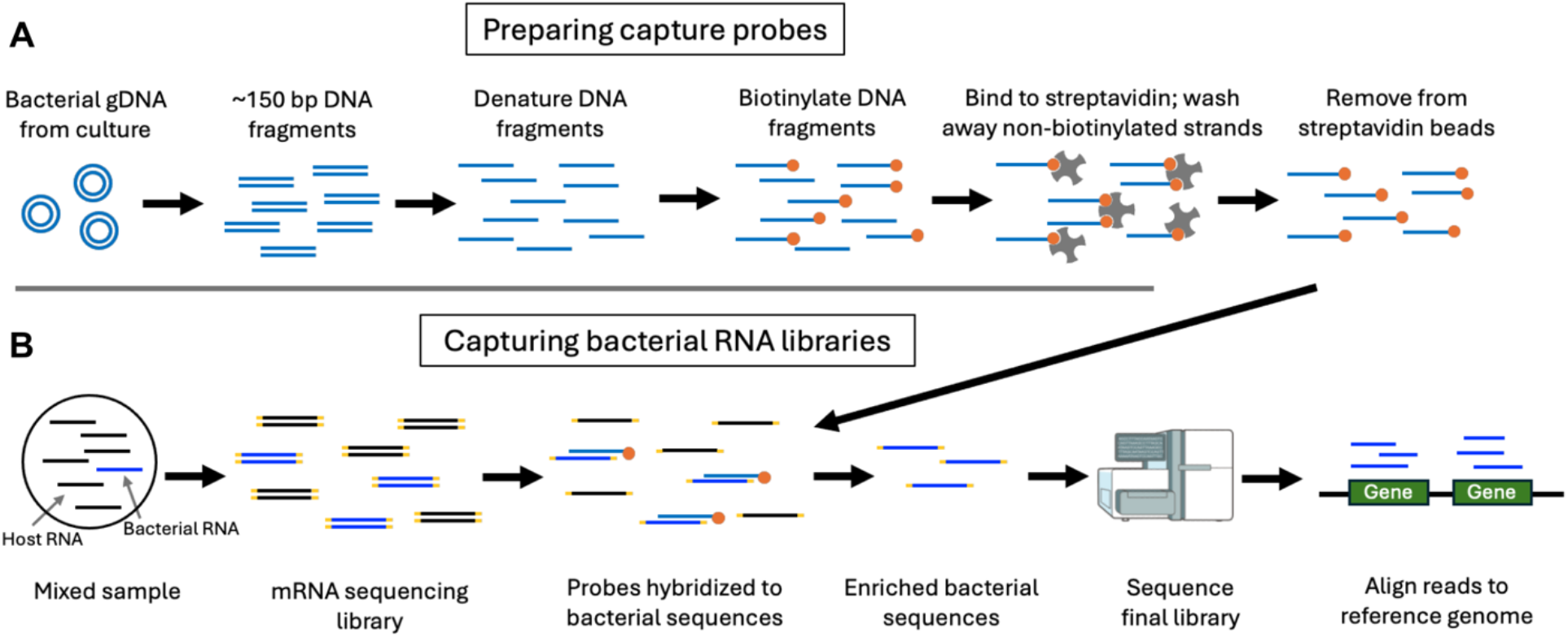
Transcript-Capture probe preparation and hybridization to sequencing libraries. A) Capture probes are generated by extracting bacterial DNA from culture, fragmenting DNA to ∼150bp, denaturing fragments, then biotinylating single stranded fragments. Non-biotinylated fragments are removed through washes with streptavidin beads. B) Bacterial RNA is captured from RNA sequencing libraries prepared from mixed samples. Probes hybridize to bacterial cDNA and are captured with streptavidin beads. Material that bound to capture probes is sequenced.

To measure the efficiency of Transcript-Capture in enriching the Mtb transcriptome, we assessed samples (n=3) prepared by spiking 1×10^6^ cells of the Mtb strain H37Ra into 1×10^6^ cells of the human monocytic cell line THP-1, which were used as a surrogate for host material. In spiked samples sequenced without Transcript-Capture, ∼60,000 reads aligned to the Mtb transcriptome out of ∼7.4 million total, comprising ∼0.8% of the library reads. After Transcript-Capture, the number of Mtb-specific reads increased to ∼15.2 million out of ∼18.8 million, averaging 80.8% of the total reads and representing a >200-fold enrichment in the number of Mtb-specific reads recovered (**Figure 2A,B**). We used the number of Mtb genes with ≥10 reads aligning as a measure of transcriptional coverage, with >80% coverage indicating robust gene representation [5,32]. Out of the 4,030 Rv genes in the annotated H37Rv genome [29,30], uncaptured samples had an average of 35% transcriptional coverage (1,450 genes with ≥10 reads aligning), after capture this increased to 99% (4,017 genes) (**Figure 2C**).

**Figure 2.**
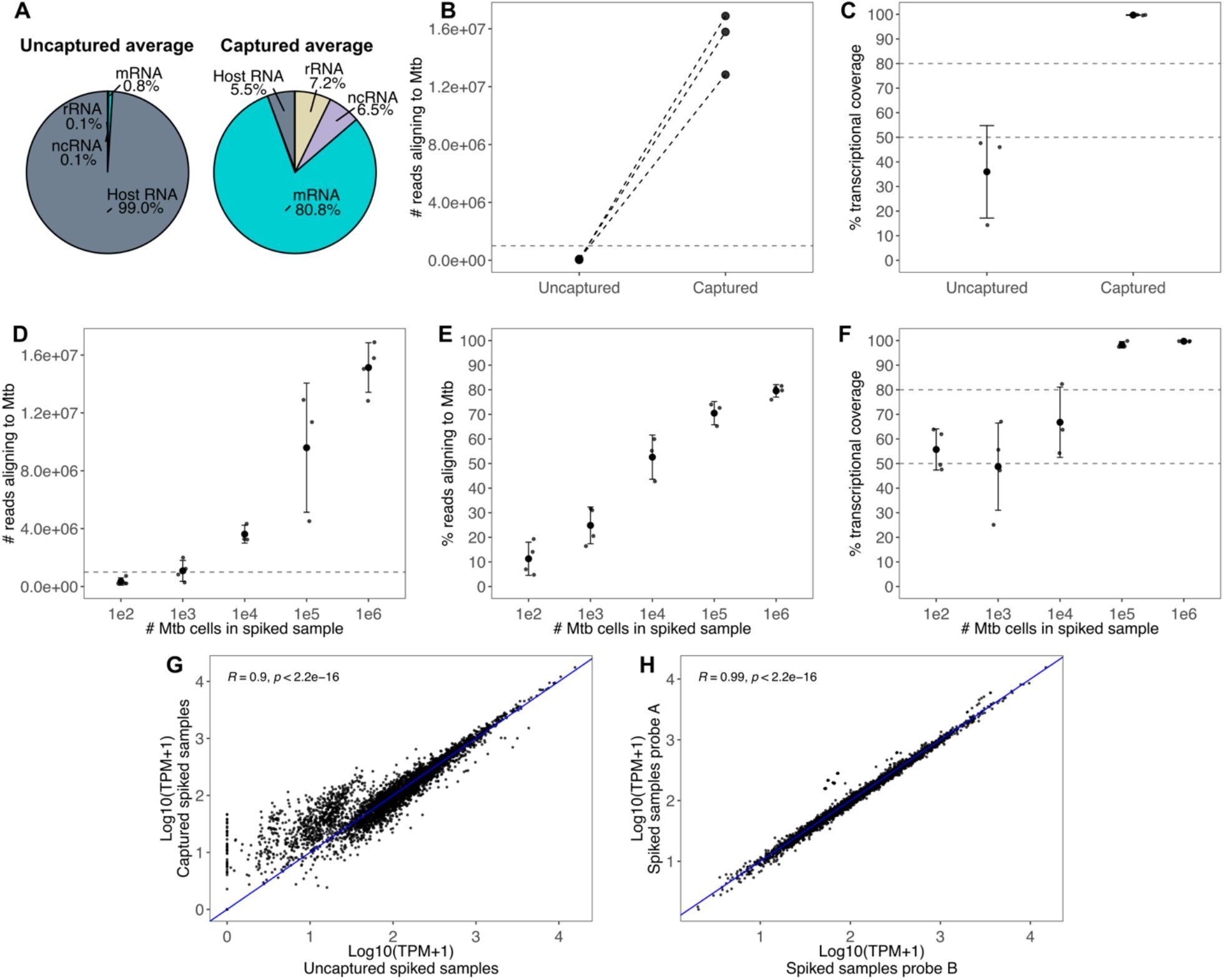
A-C: Comparison of read depth between 3 replicates of spiked samples sequenced before and after Transcript-Capture for A) percent of total reads aligning to the Mtb transcriptome; B) number reads aligning to Mtb with identical samples joined with dotted lines; C) percent of genes with at least 10 reads aligning (transcriptional coverage). D-F: Spiked samples containing 1×10^2^ to 1×10^6^ cells of Mtb of D) number reads aligning to Mtb; E) percent of total reads aligning to Mtb; F) percent of genes with at least 10 reads aligning (transcriptional coverage). Error bars indicate standard deviation. G-H: Pearson correlations of gene expression values in log_10_ transformed transcripts per million (TPM) for G) the average of 3 captured and 3 uncaptured spiked samples containing 1×10^6^ Mtb cells; H) the average of 5 spiked samples captured and sequenced twice with different probes.

To investigate the sensitivity of Transcript-Capture, we sequenced spiked samples containing between 1×10^2^ and 1×10^6^ cells of Mtb. We found that spiked samples with at least 1×10^5^ bacterial cells (n=3) had >1 million reads aligning to Mtb and >80% transcriptional coverage, demonstrating successful sequencing of the complete bacterial transcriptome (**Figure 2D-F**). As bacterial burden decreased, the percent of total reads aligning to Mtb dropped – at 1×10^2^ cells (n=4) only ∼10% of reads aligned to Mtb, and transcriptomic coverage was near 50%. While Transcript-Capture still yielded bacterial reads from these samples, only partial transcriptomes were recovered due to their lower burden. However, gene expression values normalized to transcript per million (TPM) in spiked samples with only 1×10^2^ cells remained highly correlated with spiked samples containing 1×10^6^ Mtb cells (R=0.91, p<0.05), indicating that biologically relevant information can be recovered even from 1×10^2^ bacteria and transcriptomes of partial coverage.

Probes were made from the genome of H37Rv, but our samples were spiked with a different strain. H37Ra is an avirulent Mtb with a point mutation in a single gene relative to H37Rv (*phoP*/Rv0757) [33,34]. To determine if our probes hybridize to and enrich for RNA sequences that contained SNPs, we examined the base distribution at this location. For spiked samples, the C->T SNP was observed in an average 98.2% of the reads, indicating that this approach can effectively identify SNPs present in the RNA (**Supplemental Figure 1**).

To determine if Transcript-Capture introduces expression bias, gene expression values normalized to TPM were compared between 3 spiked samples before and after capture; expression values were highly correlated (R=0.90, p<0.05) (**Figure 2G**). In uncaptured spiked samples there was a cluster of genes with low expression and >100 genes returned zero reads, likely due to the low number of reads aligning to Mtb (∼60,000). After Transcript-Capture these low-to-mid-expressed genes returned higher TPM, demonstrating that capture improves sequencing recovery of poorly expressed genes. We next assessed whether probes made from different starting cultures of H37Rv introduce expression variability. Five technical replicates of spiked samples were captured and sequenced two separate times using separate probe preparations. The samples were highly correlated with each other (R=0.99, p<0.05), indicating that different probe preparations do not introduce significant expression bias (**Figure 2H**).

### Transcript-Capture sequencing generates complete transcriptomes from biological samples

To evaluate the ability of Transcript-Capture to generate transcriptomes from a variety of sample types, we tested biological samples for which standard bacterial transcriptomics is insufficiently sensitive. We first examined samples of an *in vitro* caseum model inoculated with Mtb. Caseum mimic is made of lysed ‘foamy’ macrophages and is a surrogate for the necrotic material within TB granulomas [19]. For each sample, 1g of caseum mimic was inoculated with 1×10^8^ cells of Mtb and incubated for 7-28 days. In this model Mtb remains in a viable but mostly non-replicating state and there were 3.7×10^8^-1.3×10^9^ CFU/g of Mtb in mimic samples at the time of RNA extraction and sequencing. Transcript-Capture successfully enriched Mtb reads from this *in vitro* model containing high quantities of macrophage material; all mimic samples had >1 million reads aligning to Mtb and >80% transcriptional coverage (**Figure 3A-C**).

**Figure 3.**
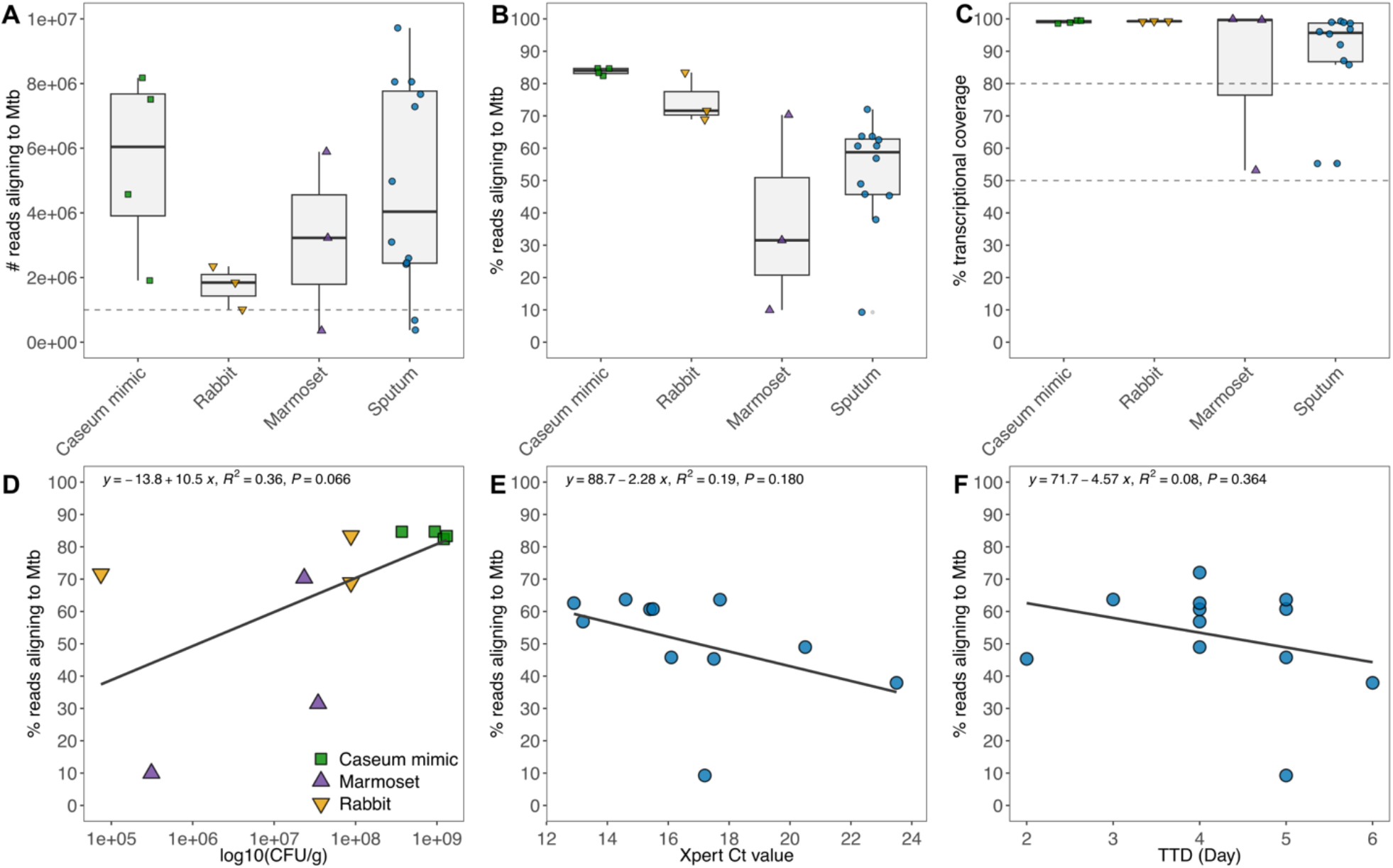
A) Number of reads aligning to Mtb, B) percent of reads aligning to Mtb, and C) number of Mtb genes with at least 10 reads aligning (transcriptional coverage) for all biological samples tested. D) CFU counts for non-clinical samples compared to percent reads aligning to Mtb. E) GeneXpert Ct values and F) Mycobacteria Growth Indicator Tube (MGIT) time to detection (TTD) values in days compared to percent reads aligning to Mtb for clinical sputum samples. Linear regression lines are shown.

Next, we performed Transcript-Capture on lesions from two animal models of TB infection: rabbit and marmoset. These models exhibit cavity disease similar to the lung lesions found in clinical pulmonary TB [22,35]. However, analyzing bacterial gene expression from tissue is challenging due to increased sample complexity and presence of host material. Rabbit samples were collected from two lesions of the same rabbit 26 weeks after aerosolized Mtb infection, and bacterial burden at time of collection ranged from 7.5×10^4^ to 8.7×10^7^ CFU/g. Sufficient reads were captured in all three rabbit samples to provide a complete bacterial transcriptome even at the lowest bacterial burden (**Figure 3A-C**). Marmoset samples were collected from a single animal 6 weeks after aerosolized Mtb infection. At time of collection, bacterial burden was ∼3×10^7^ CFU/g from two lesions and 3.1×10^5^ CFU/g from one superficially normal lung section lacking evidence of a lesion. Material from the two infected sites generated complete transcriptomes with >1 million reads aligning to Mtb and >80% transcriptional coverage (**Figure 3A-C**). The third marmoset sample, from superficially normal lung tissue, did not recover a complete transcriptome; however, despite much lower bacterial burden we recovered ∼10% of reads aligning to Mtb resulting in 50% transcriptional coverage. For all samples from our *in vitro* and *in vivo* models, linear regression revealed a trend of higher bacterial burden associated with increased percent Mtb reads recovered, but this was not significant (R^2^=0.37, p=0.06) (**Figure 3D**).

Finally, we tested Transcript-Capture on sputum samples collected from TB patients before the start of drug therapy. Each sample was from a different patient and the infecting Mtb strains were members of either Lineage 2 (L2) or Lineage 4 (L4) [36]. Sputum is another highly challenging sample type and previous NGS approaches have been limited by low read depth [18]. Of the 12 samples we sequenced, 9 had >1 million reads aligned to Mtb and >80% transcriptional coverage, demonstrating robust transcriptome recovery (**Figure 3A-C**). Two of the three remaining sputum samples still had relatively successful capture with >40% reads aligning to Mtb, and repeat sequencing to a greater depth may be able to more fully recover transcriptomes from these samples. Read percentage was compared to the clinical metrics GeneXpert qPCR Ct values and Mycobacteria Growth Indicator Tube (MGIT) time to detection (TTD) values. These are measures of bacterial burden in sputum with lower values indicating higher bacterial presence. Both metrics showed a slight but not significant negative trend with percent Mtb reads recovered for each sample (Ct: R^2^=0.18, p=0.19; TTD: R^2^=0.08, p=0.36) (**Figure 3E, F**).

### Complete transcriptomes generated by Transcript-Capture provide insight into the physiological state of bacteria *in vivo*

The transcriptomes generated by Transcript-Capture offer a glimpse into the stresses that Mtb experiences *in vivo*. We first performed a principal component analysis (PCA) comparing the captured samples with complete transcriptomes (>1 million Mtb reads and >80% transcriptional coverage) along with RNA from Mtb log phase broth culture (**Figure 4A**). All but one sputum sample clustered near Mtb broth samples. This suggests that antibiotic-naïve bacteria from different TB patient sputa share similar patterns in global gene expression both with each other and with patterns from *in vitro* broth culture. Interestingly, both marmoset and rabbit lesion samples were spread across the first principal component, suggesting higher gene expression variability in these animal lesions than was observed in sputum. The total number of differentially expressed genes (DEG) between sample types followed the trends seen in the PCA, with sputum sample gene expression most like broth and caseum mimic samples (**Figure 4B**).

**Figure 4.**
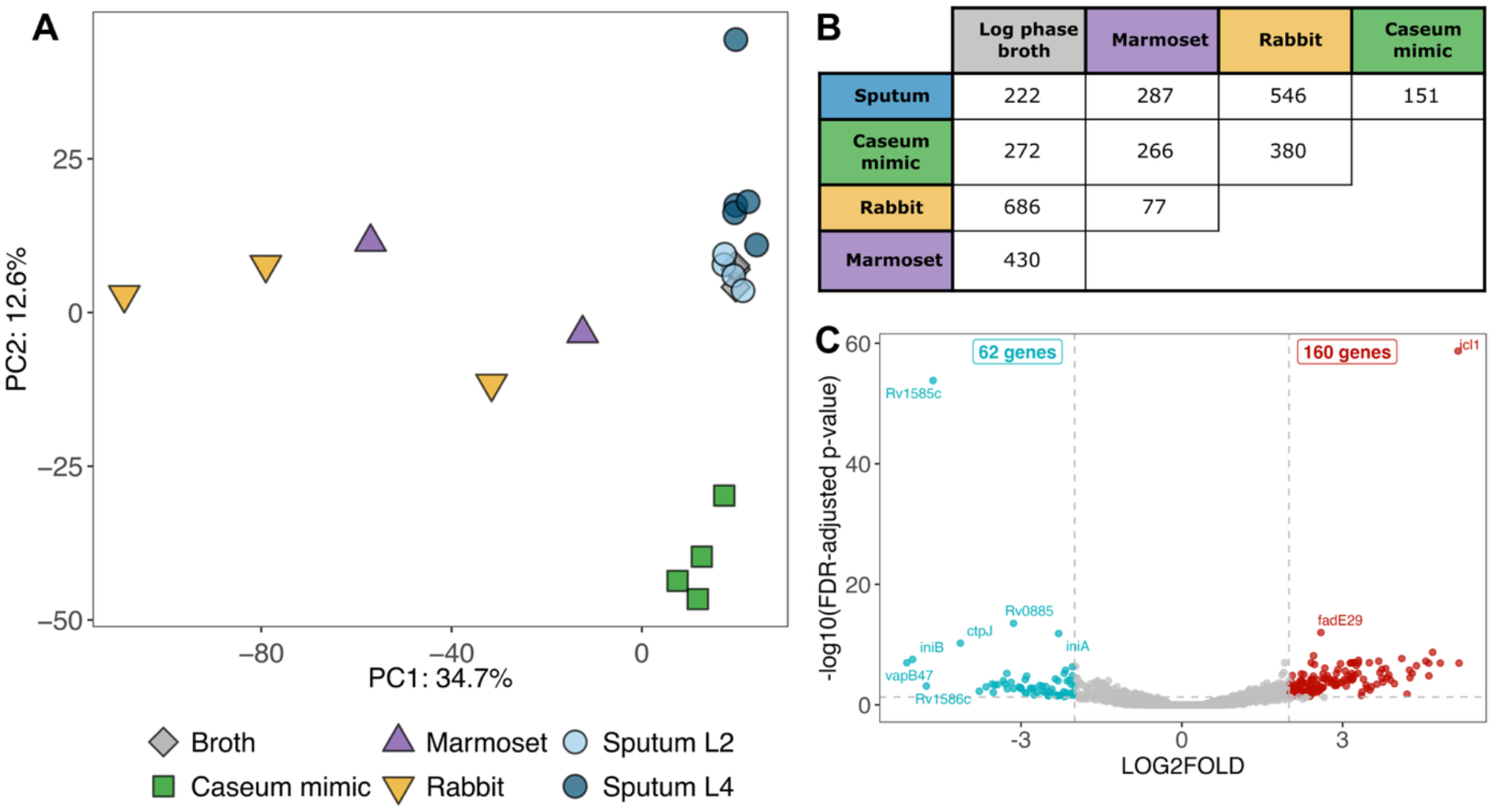
A) Principal component analysis (PCA) of all biological samples with complete transcriptomes (>1 million bacterial reads and >80% transcriptional coverage); Mtb from sputum is categorized by lineage. B) Table of significant differentially expressed gene (DEG) counts (log2fold change <-2 or >2) between each sample type; Mtb log phase broth (n=3), sputum (n=9), caseum mimic (n=4), rabbit (n=3), and marmoset (n=2). C) Volcano plot of DEG in Mtb from sputum (n=9) compared to log phase broth (n=3); red points are genes significantly upregulated in sputum and blue points are genes significantly downregulated in sputum with FDR-adjusted p-value < 0.05.

We used caseum mimic and animal lesion samples to confirm Transcript-Capture functionality across sample types but, due to small sample sizes and the difficulty in collecting additional rabbit or marmoset lesions, we did not attempt to glean biological insight from these samples. We were, however, able to generate 9 complete Mtb sputum transcriptomes (4 from L2 and 5 from L4 strains) and we performed some preliminary transcriptomic analyses on these clinical samples. Gene expression was similar between lineages, with only 45 significant DEGs between L2 and L4 samples (**Supplementary Figure S2**). These included low signal of certain genes in L2 samples that correspond with known deletions in these strains: *cas1* and *cas2* [37], *Rv0072-Rv0075* [38–40], and *PPE38* [41]. Transcript-Capture therefore allows us to tease apart lineage-specific differences in clinical samples.

We examined gene expression patterns from the average of our 9 sputum samples relative to Mtb log phase broth. To investigate transcriptional differences, we compared gene sets organized into iModulons that have been developed specifically for TB [42]. iModulons are computationally defined gene sets that represent the transcriptional regulatory networks of bacteria, and Mtb-specific iModulons split the transcriptome into 80 independent but sometimes overlapping gene sets [42,43]. The Mtb sputum transcriptome showed trends expected from bacteria surviving *in vivo*. iModulons involved in the glyoxylate and methylcitrate cycles (PrpR and BkaR) as well as lipid utilization (KsrR2 and FasR) were upregulated in sputum (**Figure 5**). Likewise, *icl1* (coding for the glyoxylate cycle enzyme isocitrate lyase) was the most significantly upregulated gene in sputum samples compared to broth (**Figure 4C**). These transcriptional profiles are consistent with the hypothesis that bacteria in the human lung are using alternative TCA cycle pathways to metabolize host lipids as a primary carbon source [18]. We also saw upregulation of iModulons IdeR and Zur, suggesting that bacteria may be in an iron- and zinc-limited environment. DevR-1 and DevR-2, iModulons encompassing the DosR regulon [26], were significantly downregulated in sputum. The iModulons Rv1828/SigH, Positive Regulation of Growth, GroEL-ES, and WhiB1 contain genes involved in regulating cell division, DNA replication, transcription, and translation [42]. These sets were similar between Mtb in sputa and log-phase broth, with only GroEL-ES iModulon showing a modest but significant decreased expression in the sputum profile (**Figure 5**).

**Figure 5.**
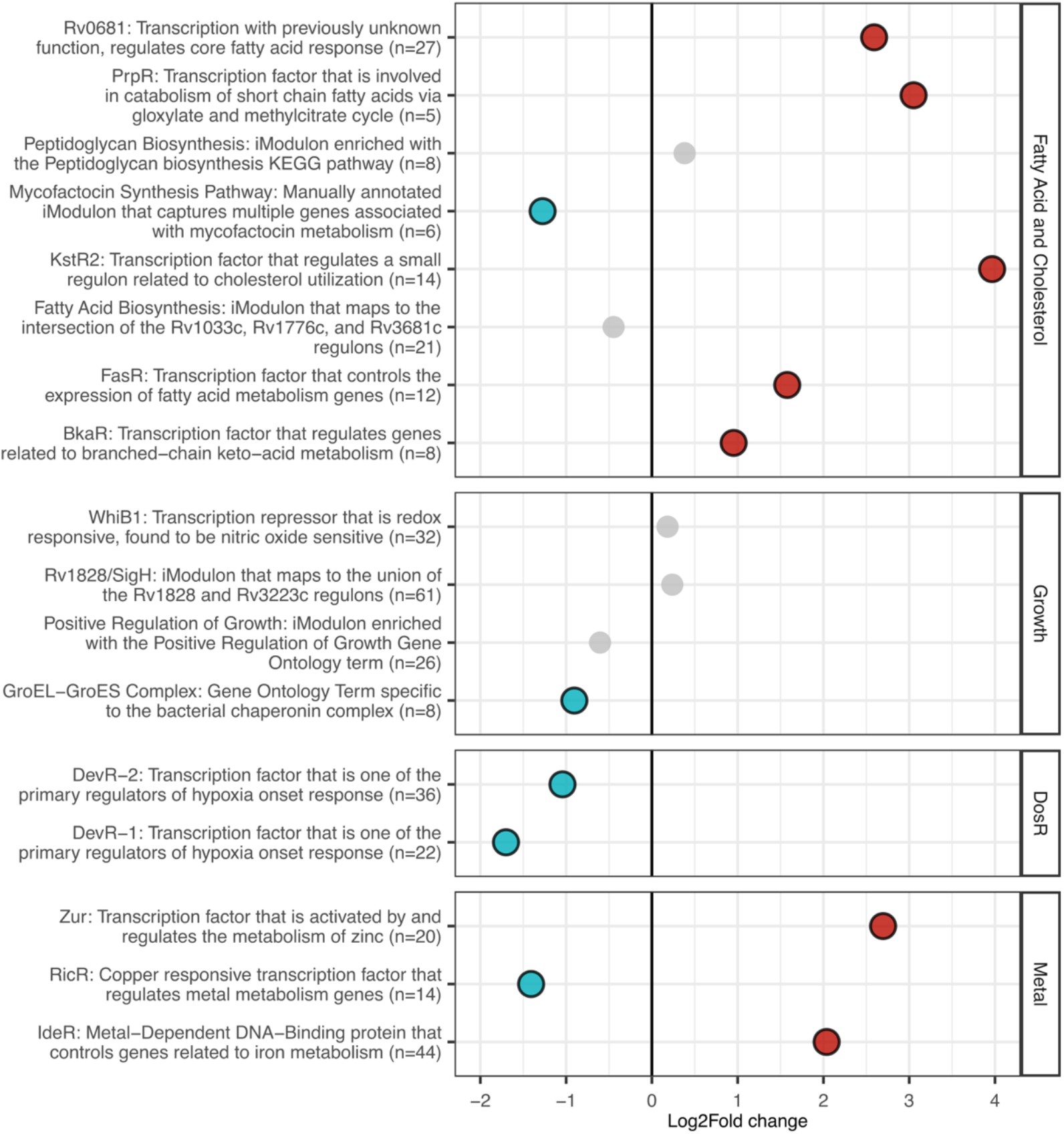
Differential expression of iModulon gene sets from Mtb in sputa (n=9) compared to Mtb log phase broth culture (n=3). Groups with significant FDR-adjusted p-values (p<0.05) are red (upregulated in sputa) or blue (downregulated in sputa).

Partial Mtb transcriptomes from antibiotic-naïve sputa have previously been generated using RT-PCR [44,45], microarrays [17,46,47], and dual RNA-seq [18]. We performed Spearman’s rank correlations comparing the average bacterial gene expression profile of TB sputum from this study with previously published expression profiles using the quantile normalization approach described in [31] (**Figure 6**). The *in vivo* transcriptome from this study showed strongest correlation (R=0.81, p<0.05) with the dual RNAseq profile from [18], minor correlation (R=0.26 and 0.34, p<0.05) with the RT-qPCR profiles from [45] and [44], and no significant correlation (R=-0.01, p=0.58) with the microarray profile from [46]. The Mtb transcriptional profiles from sputum in this current study and [18], which was analyzed using similar NGS techniques, had similar trends; both profiles revealed upregulation of genes involving host fatty acid utilization and zinc regulation, as well as a similarity in global gene expression to log phase broth culture (**Figure 4A**) ([18] Figure 3).

**Figure 6.**
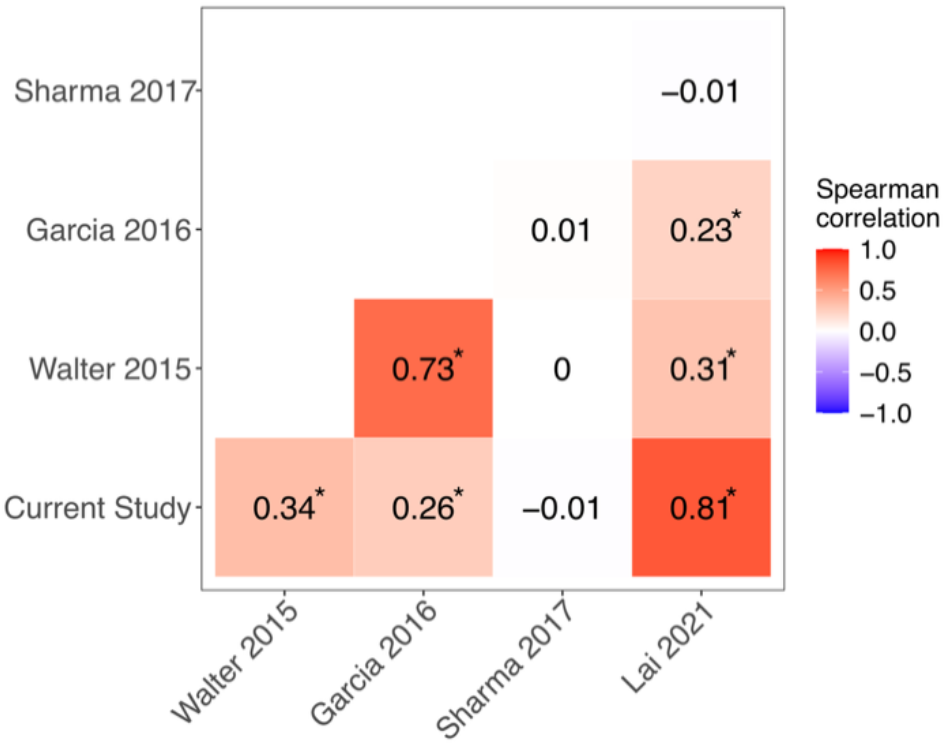
Spearman correlation of transcriptional profiles from Mtb in clinical sputa based on rank expression of Mtb genes measured in all samples (n=1,932 genes). Expression datasets are from the current study (NGS, n=9), Walter et al. (2015) (RT-PCR, n=17), Garcia et al. (2016) (RT-PCR, n=11), Sharma et al. (2017) (microarray, n=7), and Lai et al. (2021) (NGS, n=6). ^*^ indicates FDR-adjusted p-value < 0.05.

## DISCUSSION

The minimal amount of bacterial RNA and large background of host RNA present in host samples has made investigation into bacterial activity from these sources extremely challenging. Here, we report developing and deploying Transcript-Capture to enrich for bacterial RNA out of mixed samples >200-fold, including from animal models of TB infection and clinical TB sputum. Like GenCap-Seq [11], targeted enrichment using biotinylated probes generated in-house from bacterial DNA is a simple, cost-effective, and unbiased approach to sequence bacterial RNA from complex sample types such as infected host material (**Figure 1**). Although only Mtb-specific probes were used here, Transcript-Capture probes are customizable and can be used for interrogating the transcriptomes of other microbes, with the potential for wide applications across different sample types and species.

Transcript-Capture generated complete transcriptomes with >1 million bacterial reads and >80% transcriptional coverage (percent of Mtb genes with ≥10 reads aligning) from spiked samples containing at least 1×10^5^ Mtb cells (**Figure 2**). These complete transcriptomes can then be used for transcriptome-wide expression analyses. Transcriptional coverage dropped to ∼50% for samples spiked with 100-1,000 bacterial cells, but even the partial transcriptomes from samples containing 100 Mtb cells were highly correlated with those of higher bacterial burden, and 50% transcriptional coverage has been used as a benchmark in prior studies for expression analysis [18,44]. For biological samples, measures of bacterial burden were not reliable correlates of sequencing success, however a rabbit sample with just 7.5×10^4^ CFU/g generated a complete transcriptome, suggesting it is still possible to recover entire transcriptomes from lower burden samples.

Both *in vitro* and animal models of TB infection are often used as a surrogate for the human lung environment, but complete bacterial transcriptomic analyses are limited by the low ratio of bacterial to host RNA [6,48]. Here, we show that Transcript-Capture can be successfully applied to complex *in vitro* (caseum mimic) and animal (rabbit and marmoset) models to generate full Mtb transcriptomes. Bacterial RNA profiles from clinical sputa are even more difficult to generate; sputum consists of a viscous mixture of cells, mucous, and other debris often containing RNA of low quality and yield, and gene expression variability is heightened by samples from different patients infected with different clinical strains. By applying our new Transcript-Capture method to NGS libraries generated from clinical sputa, we can now directly sequence clinical samples to a depth sufficient for complete transcriptomic analysis (**Figure 3**).

Despite the inherent increase in variability expected from clinical material, we were able to discern trends in the transcriptional signature of Mtb in sputum from 9 TB infected patients. PCA analysis showed that Mtb from different patients cluster well with each other and with log phase Mtb broth culture, more so than with marmoset or rabbit lung samples (**Figure 4**). This is consistent with previous work indicating the global gene expression Mtb from sputum was similar to replicating log phase culture [18]. Expression analysis of iModulons (gene sets representing the transcriptional regulatory network of Mtb) showed that our *in vivo* Mtb expression profile shares similarities to previous descriptions of Mtb activity *in vivo*, including upregulation of genes involved in host lipid utilization and zinc limitation (**Figure 5**) [18]. Multiple lines of evidence suggest that the lung, from which sputum bacteria derive, is a hostile environment where Mtb must eat host lipids and scavenge for metals to survive [18,49–52]. Despite this potentially harsh environment, Mtb from sputa had a similar expression profile to that of log phase broth culture based on growth-related iModulons; these expression signatures suggest that Mtb in sputum may not be experiencing much growth inhibition. Although unexpected, this result is in line with the clustering in the PCA and DEG results and with the results from [18], the only other study to examine Mtb from sputa with NGS methods. Finally, we saw downregulation of the iModulons DevR-1 and DevR-2 (containing the DosR regulon). The DosR regulon is involved in the Mtb response to hypoxia and nitric oxide, and its expression increases in response to a variety of stresses [26]. The decreased expression observed here is consistent with our other indications that sputum bacilli did not experience high stress. However, this result is contrary to previous descriptions of DosR activity in sputum, which reported upregulation in Mtb from clinical sputa [44,45,47]. Documented differences in DosR response may be caused by variability of patient populations, sputum collection methods, infecting strains, disease pathology at the time of sampling, or methods of analysis. Subsequent detailed studies of sputum transcriptomics facilitated by Transcript-Capture should help to resolve this issue.

Applying Transcript-Capture to clinical sputa gives us the ability to observe a snapshot of bacterial activity directly from human patients and will be useful in future studies investigating the physiological state of bacteria surviving in the human lung. Employing this technique in patient samples with additional clinical information (for example ^18^F-FDG PET/CT scans) will allow for correlations of gene expression differences in subjects with heterogeneous pathologies. In addition, the flexibility in generating probes from different starting materials may facilitate studies with highly variable clinical isolates or mixed pathogen samples. This technique has the potential to shed new light on the stresses encountered and strategies employed by pathogens such as Mtb causing disease within their hosts and should find application in many systems where transcriptomic signal-to-noise is a limiting factor.

## ACKNOWLEDGEMENTS

The graphical abstract was created in BioRender; Lamont, E. (2025) https://BioRender.com/z2jasf2. We thank members of the Tuberculosis Imaging Program for careful conduct of the animal experiments as well as Dr. Deepti Chadalavada and the members of the Comparative Medicine Branch of NIAID for clinical monitoring and daily care of the rabbits and marmosets. We thank the Northwest Genomics Center at the University of Washington for RNA sequencing services and Erica Ryke for assistance with sequencing troubleshooting. We thank all members of the Sherman Lab including Kristin Adams for critical reading of the manuscript, and Justin Brache and José Manuel Ezquerra-Aznárez for assistance with sample generation. We thank all the TB patients who participated in the PredictTB clinical trial.

## AUTHOR CONTRIBUTIONS

EIL: Conceptualization, Formal analysis, Investigation, Methodology, Validation, Visualization, Writing – original draft, Writing – review & editing. RMJ: Conceptualization, Investigation, Methodology, Writing – original draft, Writing – review & editing. JA: Investigation, Validation, Writing – review & editing. RM: Data curation, Software. TS: Data curation, Formal analysis, investigation, Writing – review & editing. XY: Data curation. LEV: Data curation, Investigation, Project administration, Writing – review & editing. DMW: Investigation, Writing – review & editing. JW: Data curation, Project administration. SM: Conceptualization, Methodology, Writing – review & editing. RJW: Conceptualization, Funding acquisition, Writing – review & editing. CEB3: Conceptualization, Funding acquisition, Writing – review & editing. DRS: Conceptualization, Funding acquisition, Resources, Writing – original draft, Writing – review & editing.

## REFERENCES

1. Westermann AJ, Gorski SA, Vogel J. Dual RNA-seq of pathogen and host. Nat Rev Microbiol. 2012;10(9):618–630. doi:10.1038/nrmicro2852

2. Penaranda C, Hung DT. Single-Cell RNA Sequencing to Understand Host-Pathogen Interactions. ACS Infect Dis. 2019;5(3):336–344. doi:10.1021/ACSINFECDIS.8B00369

3. Dartois VA, Rubin EJ. Anti-tuberculosis treatment strategies and drug development: challenges and priorities. Nat Rev Microbiol. 2022;20(11):685–701. doi:10.1038/s41579-022-00731-y

4. Pisu D, Huang L, Grenier JK, Russell DG. Dual RNA-Seq of Mtb-Infected Macrophages In Vivo Reveals Ontologically Distinct Host-Pathogen Interactions. Cell Rep. 2020;30(2):335-350.e4. doi:10.1016/J.CELREP.2019.12.033

5. Betin V, Penaranda C, Bandyopadhyay N, et al. Hybridization-based capture of pathogen mRNA enables paired host-pathogen transcriptional analysis. Sci Rep. 2019;9(1):1–13. doi:10.1038/s41598-019-55633-6

6. Wynn EA, Dide-Agossou C, Reichlen M, et al. Transcriptional adaptation of Mycobacterium tuberculosis that survives prolonged multi-drug treatment in mice. mBio. 2023;14(6):e02363–23. doi:10.1128/mbio.02363-23

7. Peterson EJ, Bailo R, Rothchild AC, et al. Path-seq identifies an essential mycolate remodeling program for mycobacterial host adaptation. Mol Syst Biol. 2019;15(3). doi:10.15252/MSB.20188584

8. Chung M, Teigen L, Liu H, et al. Targeted enrichment outperforms other enrichment techniques and enables more multi-species RNA-Seq analyses. Sci Rep. 2018;8(1):1–12. doi:10.1038/s41598-018-31420-7

9. Donaldson GP, Chou WC, Manson AL, et al. Spatially distinct physiology of Bacteroides fragilis within the proximal colon of gnotobiotic mice. Nat Microbiol. 2020;5(5):746–756. doi:10.1038/s41564-020-0683-3

10. Long DR, Bryson-Cahn C, Waalkes A, et al. Contribution of the patient microbiome to surgical site infection and antibiotic prophylaxis failure in spine surgery. Sci Transl Med. 2024;16(742):8222. doi:10.1126/SCITRANSLMED.ADK8222

11. Hayden HS, Joshi S, Radey MC, et al. Genome Capture Sequencing Selectively Enriches Bacterial DNA and Enables Genome-Wide Measurement of Intrastrain Genetic Diversity in Human Infections. mBio. 2022;13(5). doi:10.1128/MBIO.01424-22

12. Chen RY, Via LE, Dodd LE, et al. Using biomarkers to predict TB treatment duration (Predict TB): a prospective, randomized, noninferiority, treatment shortening clinical trial. Gates Open Res. 2017;1:9. doi:10.12688/gatesopenres.12750.1

13. Ehlers S. Lazy, dynamic or minimally recrudescent? On the elusive nature and location of the mycobacterium responsible for latent tuberculosis. Infection. 2009;37(2):87–95. doi:10.1007/S15010-009-8450-7

14. World Health Organization. Global tuberculosis report 2024. 2024. Accessed November 3, 2024. https://www.who.int/publications/i/item/9789240101531

15. Dartois VA, Mizrahi V, Savic RM, Silverman JA, Hermann D, Barry CE. Strategies for shortening tuberculosis therapy. Nat Med. 2025;31(6):1765–1775. doi: 10.1038/s41591-025-03742-3

16. Colangeli R, Jedrey H, Kim S, et al. Bacterial Factors That Predict Relapse after Tuberculosis Therapy. New England Journal of Medicine. 2018;379(9):823–833. doi:10.1056/NEJMOA1715849

17. Honeyborne I, McHugh TD, Kuittinen I, et al. Profiling persistent tubercule bacilli from patient sputa during therapy predicts early drug efficacy. BMC Med. 2016;14(1):1–13. doi:10.1186/S12916-016-0609-3

18. Lai RPJ, Cortes T, Marais S, et al. Transcriptomic Characterization of Tuberculous Sputum Reveals a Host Warburg Effect and Microbial Cholesterol Catabolism. mBio. 2021;12(6). doi:10.1128/MBIO.01766-21

19. Sarathy JP, Xie M, Jones RM, et al. A Novel Tool to Identify Bactericidal Compounds against Vulnerable Targets in Drug-Tolerant M. tuberculosis found in Caseum. mBio. 2023;14(2). doi:10.1128/MBIO.00598-23

20. Via LE, Schimel D, Weiner DM, et al. Infection dynamics and response to chemotherapy in a rabbit model of tuberculosis using [18F]2-fluoro-deoxy-D-glucose positron emission tomography and computed tomography. Antimicrob Agents Chemother. 2012;56(8):4391–4402. doi:10.1128/AAC.00531-12

21. Via LE, England K, Weiner DM, et al. A Sterilizing Tuberculosis Treatment Regimen Is Associated with Faster Clearance of Bacteria in Cavitary Lesions in Marmosets. Antimicrob Agents Chemother. 2015;59(7):4181. doi:10.1128/AAC.00115-15

22. Via LE, Weiner DM, Schimel D, et al. Differential virulence and disease progression following mycobacterium tuberculosis complex infection of the common marmoset (callithrix jacchus). Infect Immun. 2013;81(8):2909–2919. doi:10.1128/IAI.00632-13

23. Malherbe ST, Chen RY, Yu X, et al. PET/CT guided tuberculosis treatment shortening: a randomized trial. medRxiv. doi:10.1101/2024.10.03.24314723, 14 October 2024, pre-print: not peer-reviewed

24. Phelan JE, O’Sullivan DM, Machado D, et al. Integrating informatics tools and portable sequencing technology for rapid detection of resistance to anti-tuberculous drugs. Genome Med. 2019;11(1):1–7. doi:10.1186/S13073-019-0650-X

25. Sherman DR, Voskuil M, Schnappinger D, Liao R, Harrell MI, Schoolnik GK. Regulation of the mycobacterium tuberculosis hypoxic response gene encoding α-crystallin. Proc Natl Acad Sci U S A. 2001;98(13):7534–7539. doi:10.1073/PNAS.121172498

26. Rustad TR, Harrell MI, Liao R, Sherman DR. The Enduring Hypoxic Response of Mycobacterium tuberculosis. PLoS One. 2008;3(1):e1502. doi:10.1371/JOURNAL.PONE.0001502

27. Culviner PH, Guegler CK, Laub MT. A Simple, Cost-Effective, and Robust Method for rRNA Depletion in RNA-Sequencing Studies. mBio. 2020;11(2). doi:10.1128/MBIO.00010-20

28. Larsen SE, Abdelaal HFM, Plumlee CR, et al. The chosen few: Mycobacterium tuberculosis isolates for IMPAc-TB. Front Immunol. 2024;15:1427510. doi:10.3389/FIMMU.2024.1427510

29. Cole ST, Brosch R, Parkhill J, et al. Deciphering the biology of mycobacterium tuberculosis from the complete genome sequence. Nature. 1998;393(6685):537–544. doi:10.1038/31159

30. Kapopoulou A, Lew JM, Cole ST. The MycoBrowser portal: A comprehensive and manually annotated resource for mycobacterial genomes. Tuberculosis. 2011;91(1):8–13. doi:10.1016/j.tube.2010.09.006

31. Coppola M, Lai RPJ, Wilkinson RJ, Ottenhoff THM. The In Vivo Transcriptomic Blueprint of Mycobacterium tuberculosis in the Lung. Front Immunol. 2021;12. doi:10.3389/FIMMU.2021.763364

32. Bhargava V, Head SR, Ordoukhanian P, Mercola M, Subramaniam S. Technical Variations in Low-Input RNA-seq Methodologies. Sci Rep. 2014;4(1):1–10. doi:10.1038/srep03678

33. Chesne-Seck ML, Barilone N, Boudou F, et al. A Point Mutation in the Two-Component Regulator PhoP-PhoR Accounts for the Absence of Polyketide-Derived Acyltrehaloses but Not That of Phthiocerol Dimycocerosates in Mycobacterium tuberculosis H37Ra. J Bacteriol. 2007;190(4):1329. doi:10.1128/JB.01465-07

34. Ryndak M, Wang S, Smith I. PhoP, a key player in Mycobacterium tuberculosis virulence. Trends Microbiol. 2008;16(11):528–534. doi:10.1016/J.TIM.2008.08.006

35. Singh AK, Gupta UD. Animal models of tuberculosis: Lesson learnt. Indian Journal of Medical Research. 2018;147(May):456–463. doi:10.4103/IJMR.IJMR_554_18

36. Gagneux S. Ecology and evolution of Mycobacterium tuberculosis. Nat Rev Microbiol. 2018;16(4):202–213. doi:10.1038/nrmicro.2018.8

37. Freidlin PJ, Nissan I, Luria A, et al. Structure and variation of CRISPR and CRISPR-flanking regions in deleted-direct repeat region Mycobacterium tuberculosis complex strains. BMC Genomics. 2017;18(1):1–14. doi:10.1186/S12864-017-3560-6

38. Chuang PC, Chen HY, Jou R. Single-Nucleotide Polymorphism in the fadD28 Gene as a Genetic Marker for East Asia Lineage Mycobacterium tuberculosis. J Clin Microbiol. 2010;48(11):4245. doi:10.1128/JCM.00970-10

39. Tsolaki AG, Gagneux S, Pym AS, et al. Genomic deletions classify the Beijing/W strains as a distinct genetic lineage of Mycobacterium tuberculosis. J Clin Microbiol. 2005;43(7):3185–3191. doi:10.1128/JCM.43.7.3185-3191.2005

40. Le Hang NT, Hijikata M, Maeda S, et al. Phenotypic and genotypic features of the Mycobacterium tuberculosis lineage 1 subgroup in central Vietnam. Sci Rep. 2021;11(1):13609. doi:10.1038/S41598-021-92984-5

41. Davies-Bolorunduro OF, Jaemsai B, Ruangchai W, Noppanamas T, Boonbangyang M, Palittapongarnpim P. Global genetic diversity of Mycobacterium tuberculosis L2.1 based on pe-ppe gene family. Infection, Genetics and Evolution. 2025;134:105802. doi:10.1016/J.MEEGID.2025.105802

42. Yoo R, Rychel K, Poudel S, et al. Machine Learning of All Mycobacterium tuberculosis H37Rv RNA-seq Data Reveals a Structured Interplay between Metabolism, Stress Response, and Infection. mSphere. 2022;7(2). doi:10.1128/MSPHERE.00033-22

43. Rychel K, Decker K, Sastry A V., Phaneuf P V., Poudel S, Palsson BO. iModulonDB: a knowledgebase of microbial transcriptional regulation derived from machine learning. Nucleic Acids Res. 2021;49(D1):D112–D120. doi:10.1093/NAR/GKAA810

44. Walter ND, Dolganov GM, Garcia BJ, et al. Transcriptional Adaptation of Drug-tolerant Mycobacterium tuberculosis During Treatment of Human Tuberculosis. J Infect Dis. 2015;212(6):990–998. doi:10.1093/INFDIS/JIV149

45. Garcia BJ, Loxton AG, Dolganov GM, et al. Sputum is a surrogate for bronchoalveolar lavage for monitoring Mycobacterium tuberculosis transcriptional profiles in TB patients. Tuberculosis. 2016;100:89–94. doi:10.1016/J.TUBE.2016.07.004

46. Sharma S, Ryndak MB, Aggarwal AN, et al. Transcriptome analysis of mycobacteria in sputum samples of pulmonary tuberculosis patients. PLoS One. 2017;12(3):e0173508. doi:10.1371/JOURNAL.PONE.0173508

47. Garton NJ, Waddell SJ, Sherratt AL, et al. Cytological and Transcript Analyses Reveal Fat and Lazy Persister-Like Bacilli in Tuberculous Sputum. PLoS Med. 2008;5(4):e75. doi:10.1371/JOURNAL.PMED.0050075

48. Westermann AJ, Vogel J. Cross-species RNA-seq for deciphering host–microbe interactions. Nat Rev Genet. 2021;22(6):361–378. doi:10.1038/S41576-021-00326-Y

49. Daniel J, Maamar H, Deb C, Sirakova TD, Kolattukudy PE. Mycobacterium tuberculosis Uses Host Triacylglycerol to Accumulate Lipid Droplets and Acquires a Dormancy-Like Phenotype in Lipid-Loaded Macrophages. PLoS Pathog. 2011;7(6):e1002093. doi:10.1371/JOURNAL.PPAT.1002093

50. Muñoz-Elías EJ, McKinney JD. Mycobacterium tuberculosis isocitrate lyases 1 and 2 are jointly required for in vivo growth and virulence. Nat Med. 2005;11(6):638–644. doi:10.1038/nm1252

51. Rodriguez GM, Sharma N, Biswas A, Sharma N. The Iron Response of Mycobacterium tuberculosis and Its Implications for Tuberculosis Pathogenesis and Novel Therapeutics. Front Cell Infect Microbiol. 2022;12:876667. doi:10.3389/FCIMB.2022.876667

52. Dow A, Sule P, Odonnell TJ, et al. Zinc limitation triggers anticipatory adaptations in Mycobacterium tuberculosis. PLoS Pathog. 2021;17(5):e1009570. doi:10.1371/JOURNAL.PPAT.1009570

