## SupplementaryMaterials for "Transcript-Capture sequencing enriches mRNA of *Mycobacterium tuberculosis* from host samples"

Supplementary Methods

#### Probe Generation

**1. Fragment genomic DNA**

a. Start with gDNA that has been extracted in a way to prevent shearing (no bead beating)

1.1 Using a Covaris, shear DNA to an average of 150bp fragments. Maximum amount of DNA per Covaris tube is 5µg. Always use 130µL volume in the 130µL milliTUBE to keep from shattering.

(Starting with 10x5µg will generate roughly 2-3µg of probe)

**Treatment conditions below:**

| Treatment Time | Setpoint Temp, °C | Min Temp, °C | Max Temp, °C | Peak Power | Cyclers/Burst | Duty Factor |
|---|---|---|---|---|---|---|
| 585 sec | 20.0 | 18.0 | 75.0 | 50 | 200 | 15% |

1.2 Use LoBind tubes/plates from here on.

**2 Clean up using Ampure XP beads**

** Bring Ampure XP (Beckman Coulter #A63882) beads to room temperature*

2.1 Aliquot 50µL sheared DNA per well.
2.2. Add 150µL (3X) Ampure beads to each well. Pipette to mix thoroughly.
2.3. Incubate at room temperature for 5 minutes.
2.4. Place on magnet to clear beads ~5 minutes.
2.5. Discard supernatant.
2.6. Add 200µL 80% ethanol and incubate at room temp for 30 sec.
2.7. Discard supernatant.
2.8. Repeat ethanol wash once.
2.9. Discard supernatant and remove residual ethanol. Allow to air dry for 5 minutes on magnet, remove excess ethanol with smaller tip.
2.10. Remove from magnet and resuspend beads in 40µL DEPC H2O. Mix well. Incubate for 5 min at room temp.
2.11. Clear beads on magnet for ~2 minutes.
2.12. Transfer all samples to same 5mL lo-bind tube.

**3 Quantitate using Qubit**

*Expected dsDNA concentration: 10-30ng/µL (max has been 50ng/µL)*

3.1. Aliquot 500ng of DNA into wells of a lo-bind PCR plate.

*****Stopping Place: Samples can be stored overnight at 4C.**

**4 Dephosphorylation the ends of sheared DNA**

4.1. Make a master mix of dephosphorylation reagents and add to sheared DNA for a total volume of 50µL per reaction:

| COMPONENT | VOLUME PER SAMPLE |
|---|---|
| Cutsmart Buffer (10X) | 5 µL |
| rSAP (NEB #M0371S) | 2.5 µL |
| Sheared + cleaned DNA (500ng) | variable |
| H2O to 50µL | variable |
| Total volume | 50 µL |

Add 7.5µL MM to each well of 500ng DNA.

4.2. Incubate at 37°C for 2 hours followed by 65°C for 15 minutes.

5 **Denaturation**

5.1. Heat samples in thermocycler at 100C for 10min with lid set to 105C.

5.2. Immediately flash freeze plate in dry ice + EtOH bath to prevent renaturation (alternatively can immediately place on ice for >5min).

6 **Biotinylation**

6.1. To the reaction from step 5.2 add reagents for Biotinylation

| COMPONENT | VOLUME PER SAMPLE |
|---|---|
| 2.4mM CoCl2 | 5 µL |
| 10µM Biotin-11-ddATP (1:100 dilution in H2O) (Revvity NEL548001EA) | 2.5 µL |
| Terminal Transferase (NEB M0315L) | 1 µL |
| Dephosphorylated DNA | 50 |
| Total volume | 58.5 µL |

6.2. Incubate for 1.5 hours at 37°C followed by heat inactivation at 70°C for 10 minutes.

6.3. Leave at 4°C overnight in the PCR block or in a refrigerator, or continue to Step 6.

*****Stopping Place: Samples can be stored overnight at 4°C before or after Monarch Kit Clean up.**

6.4. Pool all samples into one 5mL lo-bind tube.

6.5. Quantitate using Qubit ssDNA HS kit.

*Expected ssDNA concentration: 10-30ng/µL*

7 **Remove unbound biotin with Monarch PCR & DNA Cleanup Kit (NEB T1030, ssDNA protocol)**

*Turn on heat block to 50°C and heat DEPC H2O*

7.1 Aliquot DNA into <5µg samples in separate 5mL tubes. Samples should be 300-500µL.

*** We have been using 400µL sample for each column regardless of concentration*

7.2. Add 100µL DNA Cleanup Binding buffer for every 50µL sample.

For a 400µL sample, add 800µL Binding buffer

Mix by pipetting, **do not vortex!**

7.3. Add 300µL 95% EtOH for every 50µL sample.

For a 400µL sample, add 2400µL 95% EtOH

Mix by pipetting, **do not vortex!**

7.4. Insert column into collection tube and load 800µL of sample onto column. Spin for 1 minute @ 16,000xg (~13,000 rpm) then discard flow-through.

7.5. Repeat for each sample by adding more sample to the column without exceeding 800uL. Repeat until all sample has gone through the column.

7.6. Re-insert column into collection tube. Add 500µL of DNA Wash Buffer and spin for 1 min. Discard flow-through.

7.7. Repeat step 7.6 two more times. Total of 3 washes.

7.8. After discarding flow-through from the final wash, reinsert column into collection tube and spin again to make sure membrane is dry.

7.9. Transfer column to a clean 1.5mL microfuge tube.

7.10. Add 50µL of warm H2O to the center of the matrix. Wait for 1 min, then spin for 1 minute to elute DNA.

7.11. Add another 50µL H2O to the center of the matrix. Wait for 1min, then spin for 1 minute to elute DNA. Final volume is 100µL.

8 **Purify with MyOne Streptavidin C1 to eliminate non-biotinylated strands**

**** Turn on heat block to 65C for formamide disassociation step.*

8.1. Wash Dynabeads MyOne Streptavidin C1 (Invitrogen 65002):

8.1.1. Aliquot 50µL Dynabeads per reaction into a 1.5mL LoBind tube. Maximum 6 reactions per tube.

8.1.2. Place beads on magnet until clear, ~2 minutes.

8.1.3. Discard supernatant.

8.1.4. Add an equal volume (or at least 1mL) of low salt 1X Binding & Wash buffer and resuspend.

8.1.5. Place beads on magnet until clear (~2 min), discard supernatant.

8.1.6. Wash 3 times as follows:

a. Remove tube from magnet and resuspend beads in volume of low salt 1X Binding & Wash buffer equal to the initial volume of beads taken.

b. Pipet to resuspend the beads.

c. Clear beads on magnet (~1 min).

d. Discard supernatant.

8.1.7. After 3 washes, resuspend beads in low salt **2X** Binding & Wash Buffer at twice the original volume per reaction.

8.1.8. Aliquot 100µL of resuspended beads into separate 1.5mL LoBind tubes.

8.2. Add the 100µL DNA sample from step 7.11 to the washed beads.

8.2.1. Gently pipet to resuspend the beads.

8.2.2. Incubate at RT for 15 minutes (not on the magnet). Tap or vortex tube to mix beads 3-4 times during incubation.

8.3. Put tube on magnet (~2 min).

8.4. Resuspend beads in 200µL low salt 1X Binding & Wash Buffer. Place on magnet ~2min. Remove supernatant. Binding of DNA is now complete.

8.5. Resuspend beads in 100µL 0.1X SSC. Place in magnet ~2min. Remove supernatant.

8.6. Resuspend beads in 50µL formamide mix (see recipe selection).

9 **Dissociate biotinylated DNA from beads using formamide**

9.1. Incubate sample from previous step at 65°C for 5 minutes.

9.2. Place on magnet to clear beads ~2 minutes.

9.3. Transfer supernatant (50µL) to clean tube.

10 **Remove formamide with Monarch PCR & DNA Cleanup Kit using ssDNA Method**

*Turn on heat block to 50°C and heat Elution buffer*

10.1 A starting sample volume of 50µL is recommended. For smaller samples, nuclease-free water can be used to adjust the volume.

10.2 Add 100µL DNA Cleanup Binding Buffer to the 50µL sample.

10.3 Add 300µL ethanol (≥ 95%). Mix well by pipetting up and down or flicking the tube. **Do not vortex!**

10.4 Insert column into collection tube, load sample onto column and close the cap. Spin for 1 minute @16,000xg, then discard flow-through.

10.5 Re-insert column into collection tube. Add 500µL DNA Wash Buffer and spin for 1 minute @16,000xg. Discard flow-through.

10.6 Repeat Step 10.5.

10.7 Transfer column to a new LoBind 1.5 ml microfuge tube. Use care to ensure that the tip of the column does not contact the flow-through. If in doubt, re-spin for 1 minute to ensure traces of salt and ethanol are not carried over to the next step.

10.8. Add 11µL of warm Elution Buffer to the center of the matrix. Wait for 1 min, then spin for 1 minute to elute DNA.

10.9. Add another 11µL Elution Buffer to the center of the matrix. Wait for 1min, then spin for 1 minute to elute DNA.

11 **QC**

11.1. Quantitate with ssDNA HS qubit.

*Expected ssDNA concentration: 15-25ng/μL*

11.2. Check biotinylation with dot blot (1μL).

**<u>Recipes:</u>**

**5M NaCl**

14.61g NaCl to 50 ml water - heat gently to dissolve

**Low Salt 2X Binding and Wash Buffer Recipe (300mM NaCl)** (makes 50 ml)

10 mM Tris-HCl
1 mM EDTA
300mM NaCl

To make 50mL

- 3 mL of 5M NaCl
- 0.5mL of 1M Tris-HCl
- 0.1mL of 0.5M EDTA
- 46.4mL ultrapure water
- Filter through a 0.2μm syringe filter to sterilize

**1X Binding and Wash Buffer**

Dilute 2X buffer with equal volume water

**0.1X SSC**

To make 50mL

- 49.75mL H2O
- 250μL 20X SSC

**Formamide** = 950ul formamide + 20ul (0.5M) EDTA + 30ul water. Aliquot and store in -20. Avoid freeze thaws.

### Dot Blot

**Materials**

- Nitrocellulose membrane, 0.2µm (Thermo Scientific #88024)
- Blocking buffer:
  - 2% BSA: 20g BSA (Sigma #A5611) in 1L 1X TBST
- Wash buffer (1X TBST):
  - 1X TBST: TBST = Tris buffered saline with 0.1% Tween20
    50mL 10X TBS
    450mL MilliQ H2O
    0.5mL Tween20
  - To make 10X TBS:
    12g Tris base
    44g NaCl
    400mL MilliQ $H_2O$
    Adjust to pH 7.6 with 12N HCL
    Adjust to final volume 500mL with MilliQ $H_2O$
- Streptavidin-AP Conjugate options:
  - Streptavidin-AP conjugate (Roche #11089161001)
- NBT/BCIP Substrate solution:
  - BCIP/NBT Liquid Substrate System (Sigma #B1911-100ML)

**Protocol**

1. Prepare ssDNA samples
   a. Positive control: Prepare dilutions of biotinylated DNA with known molarity
2. Load nitrocellulose membrane
   a. Apply 1µL dots of samples and positive controls to the membrane
3. Immediately dry the dot blot
   a. Use a UV crosslinker according to the manufacturer's instructions to fix the DNA to the membrane
4. Blocking
   a. Immerse the membrane in blocking buffer and incubate for 1hr at RT with gentle shaking to block non-specific binding sites
   b. Wash the membrane 3 times with wash buffer, 5-10 min per wash, with gentle shaking
5. Incubate with Streptavidin-AP conjugate
   a. Dilute Streptavidin-AP conjugate in blocking buffer
      i. Dilution varies, typically 1:1000 to 1:5000 dilution. (Have used 1:2000)
   b. Immerse the membrane in diluted Streptavidin-AP conjugate (~10-20mL) and incubate for 1hr at RT with gentle shaking
6. Washing
   a. Wash the membrane 3 times with wash buffer, 10 min per wash, with gentle shaking

7. Detection (Colorgenic)
    a. Detection with Sigma protocol:
        i. Incubate the membrane in the substrate mixture for 10-30 min until color develops
        ii. Stop the reaction by washing the membrane in several changes of distilled H2O
        iii. Air dry the membrane and store in the dark

### Transcript Capture

**Thermal Cycler Settings:**
<u>HYB program (lid set at 100°C)</u>
95°C – 10 min
65°C – 16 hrs*
65°C – Hold
*Duration of hybridization should be kept consistent for all samples within a project. For GC-rich or small panels (<1000 probes), longer hybridization times (up to 16hr) may improve performance.
<u>WASH program (lid set at 70°C*)</u>
It is critical to reduce the lid temperature to 70°C for the WASH program.
65°C - Hold

**Day 1**

1. **Hybridization reaction**

1.1 Aliquot 500ng of each sequencing library into individual 1.5mL LoBind tubes.
* Libraries can be pooled into the same tube. We have pooled up to 5 libraries (500x5ng total).
* Make sure there is at least 500ng in each tube. Pool libraries if necessary to reach this number.

1.2 Create the Blocker Master Mix, add an additional sample for error:

| **Blocker Master Mix components** | **Volume per reaction (ul)** |
|---|---|
| Human Cot DNA | 5 |
| xGen Blocking Oligos | 2 |

1.3 Vortex to mix well and add to samples.
1.4 Dry down the mixture in a SpeedVac (65°C for 20 min or more depending on the library volume), ensure all the contents are dry. Create the master mix and buffers below while you wait for your samples to dry.

*****Stopping Place. Store samples at room temperature overnight, or -20°C for longer.**

1.5 Bring contents of the xGen Hybridization and Wash Kit (IDT 1080584) to room temperature. Inspect the tube of 2X Hybridization Buffer for precipitate. If crystals are present, heat the tube at 65°C, shaking intermittently.

1.6 Create the Hybridization Master Mix:

| **Hybridization Master Mix component** | **Volume per reaction (ul)** |
|---|---|
| xGen 2X Hybridization Buffer | 8.5 |
| xGen Hybridization Buffer Enhancer | 2.7 |
| Custom probes* + H2O | 5.8* |

* Custom probe concentration: Variable, this is what has worked in the past:

| Number of pooled libraries | ng probe per reaction |
|---|---|
| 1 | 20-25 |
| 2-3 | 40-50 |
| 4-5 | 50 |

Combine the desired concentration of probe and NF $H_2O$ to make 5.8µL total

1.7 Vortex to mix, add 17µL to each well of the tubes containing dried DNA.
1.8 Transfer contents from the tubes to individual wells of a 96 well Lo bind plate.
1.9 Securely seal the plate with a Microseal B seal.
1.10 Incubate at least 5 min at room temperature.
1.11 Vortex the samples, making sure that they are completely mixed.
1.12 Briefly centrifuge the samples.
1.13 Place the plate on the thermal cycler and start the HYB program.
1.14 Prepare the buffers if you have not already done so.

#### 2. Prepare Buffers

2.1 Dilute the following xGen buffers to create 1X working solutions as follows, multiplying by the required number of samples and adding 10% extra:

| Component | Nuclease-Free Water (µL) | Buffer (µL) | Total (µL) |
|---|---|---|---|
| xGen 2X Bead Wash Buffer | 150 | 150 | 300 |
| xGen 10X Wash Buffer 1* | 225 | 25 | 250 |
| xGen 10X Wash Buffer 2 | 135 | 15 | 150 |
| xGen 10X Wash Buffer 3 | 135 | 15 | 150 |
| xGen 10X Stringent Wash Buffer | 270 | 30 | 300 |

***Note:** If Wash Buffer 1 is cloudy, heat the bottle in a 65°C water bath to allow resuspension.

2.2 Swirl to mix. Do not vortex.
2.3 The 1X working solutions are stable at room temperature (15–25°C) for up to 4 weeks.
2.4 Use a fresh PCR plate. For 24 samples, as an example, aliquot and label the plate as follows:
- Rows 1–2: 110 µL of Wash buffer 1
- Rows 3-4: 160 µL of Stringent Wash Buffer
- Rows 5-6: 160 µL of Stringent Wash Buffer

2.5 Do not discard the remaining Wash Buffer 1. The remaining buffer is needed to perform the Room temperature washes later in the protocol.
2.6 Seal the buffer plate and set aside.

2.7 In a LoBind tube, make the Bead Resuspension Mix. Multiply by the number of samples and add a 10% extra.

| Bead Resuspension Mix component | Volume per reaction (ul) |
|---|---|
| xGen 2X Hybridization Buffer | 8.5 |
| xGen Hybridization Buffer Enhancer | 2.7 |
| Nuclease-Free Water | 5.8 |

**Day 2**

3. **Wash Streptavidin beads**

***Remove the Dynabeads M-270 Streptavidin (Fisher 65-305) beads from storage at 4°C and equilibrate the beads at room temperature at least 30 min before performing the washes.***

3.1 Mix the beads thoroughly by vortexing for 15 sec.
3.2 Add 50µL of Streptavidin beads to a new PCR plate, filling a well for every sample being captured.
3.3 Add 100µL of Bead Wash Buffer from Prepare buffers, step 1 to each well, then gently pipet the mix 10 times.
3.4 Place the plate containing beads on a magnet and allow the beads to fully separate from the supernatant (approximately 1 min).
3.5 Remove and discard the clear supernatant, ensuring that the beads remain in the well.
3.6 Remove the plate containing beads from the magnet.
3.7 Perform the following wash:
   - Add 100µL of Bead Wash Buffer to each well containing beads, then gently pipet the mix 10 times.
   - Place the plate on the magnet for approximately 1 min, allowing beads to fully separate from the supernatant.
   - Carefully remove and discard the clear supernatant.

3.8 Perform an additional wash by repeating step 7 (above) for a total of 2 washes.
3.9 Resuspend the beads in 17µL of Bead Resuspension Mix from Prepare buffers, step 6.
3.10 Mix thoroughly to ensure the beads are not left to dry in the well. If needed, briefly centrifuge the plate containing beads at 25 x *g* (400 rpm).

4. **Perform bead capture**

***If any of the sample accidentally splashes onto the plate seal while vortexing in Perform bead capture, briefly and gently centrifuge the plate (10 sec at 25 x g).***

4.1 Start the WASH program in the second thermal cycler to start warming the buffer plate prepared in Prepare buffers, step 2. Make sure the lid temperature is set to 70°C for the WASH program
4.2 The buffer plate needs to warm up for at least 15 min. Recommend starting incubation at the same time as the bead capture.

4.3 After the hybridization incubation is complete, remove the sample plate from the thermal cycler.
4.4 Once the sample plate has been removed from the instrument, stop the HYB program.
4.5 Immediately after the HYB program is complete, start the WASH program. At this point, both thermal cyclers should be running the WASH program.
4.6 Using a multichannel pipette and fresh filter tips, transfer the fully homogenized beads to the samples and gently pipette until fully mixed.
4.7 Securely seal the sample plate.
4.8 Place the sample plate in the thermal cycler for 45 min. During incubation, remove the plate every 10–12 min to quickly and gently vortex.

***It is safe to place the sample plate in the thermal cycler before the lid temperature has fully cooled to 70°C when starting the incubation.***

**<u>Perform washes</u>**

Always keep the buffer plate on the thermal cycler during washes. Make sure to reseal the buffer plate in between washes. When performing the heated washes, keep the buffer plate on the thermal cycler to maintain its set temperature.

<u>Heated washes</u>

1. After 45 min, remove the sample plate from the thermal cycler.
2. With the buffer plate remaining in the thermal cycler, transfer 100µL of heated Wash Buffer 1 to each sample and pipet the mix 10 times, being careful to minimize bubble formation.
3. Reseal the buffer plate, then close the lid.
4. Place the sample plate on the magnet for 1 min. Remove the supernatant and discard.
5. Remove the sample plate from the magnet, then add 150ul of heated Stringent Wash Buffer to each well containing a sample. Reseal the buffer plate, then close the lid.
6. Pipet the mix 10 times, being careful to minimize bubble formation. Always use fresh pipette tips for each well.
7. Securely seal the sample plate, then incubate for 5 min in the thermal cycler.
8. Place the sample plate on the magnet for 1 min, then remove the supernatant and discard.
9. Remove the sample plate from the magnet, then add 150µL of heated Stringent Wash Buffer from the buffer plate to the sample plate.
10. Pipet the mix 10 times, being careful to minimize bubble formation. Securely seal the sample plate, then incubate for 5 min on the thermal cycler.
11. Place the sample plate on the magnet for 1 min.

<u>Room temperature washes</u>

To ensure that the beads remain fully resuspended, vigorously mix the samples during the room temperature washes.

1. Remove supernatant. Add 150μL of Wash Buffer 1.
2. Securely seal the sample plate with a fresh seal, then vortex at full-speed thoroughly, until fully resuspended. It is critical to use a new seal at this step to avoid the risk of contamination because there will be some bead splash on the seal.
3. Incubate for 2 min while alternating between vortexing for 30 sec and resting for 30 sec, to ensure the mixture remains homogenous.
4. Centrifuge the sample plate for 5 sec at 25 x *g* to avoid well-to-well contamination.
5. Place the sample plate on the magnet for 1 min, then remove and discard the seal.
6. Remove the supernatant, then remove the sample plate from the magnet.
7. Add 150μL of Wash Buffer 2, then securely seal the sample plate with a fresh seal and vortex thoroughly until fully resuspended. *Beads tend to stick to wells after Wash 2.
8. Incubate for 2 min while alternating between vortexing for 30 sec and resting for 30 sec, to ensure the mixture remains homogenous.
9. After the incubation, briefly centrifuge the sample plate (5 sec at 25 x *g*).
10. After centrifuging, place the sample plate on the magnet for 1 min, then remove and discard the seal.
11. Remove the supernatant, then remove the sample plate from the magnet.
12. Add 150μL of Wash Buffer 3, then securely seal the sample plate with a fresh seal and vortex thoroughly until fully resuspended.
13. Incubate for 2 min while alternating between vortexing for 30 sec and resting for 30 sec, to ensure the mixture remains homogenous.
14. After the incubation, briefly centrifuge the sample plate (5 sec at 25 x *g*).
15. After centrifuging, place the sample plate on the magnet for 1 min, then remove and discard the seal.
16. Remove the supernatant.
17. With the sample plate still on the magnet, use fresh pipette tips to ensure that all residual Wash Buffer 3 has been removed, then remove the plate from the magnet.
18. Add 20μL of Nuclease-Free Water to each capture.
19. Pipet the mix 10 times to resuspend any beads stuck to the side of the well.
20. Do not discard the beads. Use the entire 20μL of resuspended beads with captured DNA in Perform post-capture PCR.

**<u>Perform post-capture PCR</u>**

1. In a tube, prepare the Amplification Reaction Mix, multiplied by the number of samples on the plate and adding 10% overfill, as follows:

| **Amplification Reaction Mix component** | **Volume (μL)** |
|---|---|
| 2X KAPA HiFi HotStart ReadyMix | 25 |

| | |
|---|---|
| 10 µM Illumina P5 primer | 2.5 |
| 10 µM Illumina P7 primer | 2.5 |
| Beads with captured DNA from step. 20 | 20 |

P5 primer: AATGATACGGCGACCACCGA
P7 primer: CAAGCAGAAGACGGCATACGA

1. Mix for a final reaction volume of 50µL.
2. Securely seal the sample plate, then pipette 10x to thoroughly mix the reaction.
3. Briefly centrifuge the plate.
4. Place the plate in a thermal cycler, and run the following program with the lid temperature set to 105°C:

| **Step** | **Number of cycles** | **Temperature (°C)** | **Time** |
|---|---|---|---|
| Polymerase activation | 1 | 98 | 45 sec |
| Denaturation | Variable (15-20) | 98 | 15 sec |
| Annealing | | 60 | 30 sec |
| Extension | | 72 | 30 sec |
| Final extension | 1 | 72 | 1 min |
| Hold | 1 | 4 | Hold |

***Stopping Place**. Amplified captures may be stored at 4°C overnight.

**Day 3**
**Purify post-capture PCR fragments**

Ensure Ampure XP beads (Beckman Coulter Life Sciences) have been equilibrated to room temperature before proceeding.

1. Prepare 250µL of fresh 80% ethanol per sample, multiplied by the number of samples with 10% extra.
2. Add 75µL (1.5X volume) of Ampure XP beads to each amplified capture.
3. After adding the beads, pipet the mix thoroughly and incubate for 5–10 min.
4. Place the plate on the magnet until the supernatant is clear (2–5 min).
5. Remove the supernatant without disturbing the beads.
6. While keeping the plate on the magnet, add 125µL of 80% ethanol, then incubate for 1 min.
7. Remove the ethanol, then repeat another ethanol wash.
8. Allow the beads to air dry for 1–3 min. Do not overdry the beads.
9. Remove the sample plate from the magnet and elute in 22µL of Buffer EB, or equivalent (10 mM Tris-Cl, pH 8.5). Mix thoroughly.
10. Incubate for 5 min at room temperature.
11. Place the plate on a magnet until supernatant is clear (1–2 min).

12. Transfer 20µL of eluate to a fresh plate, making sure that no beads are carried over.

*Purified PCR fragments may be stored at -20°C for up to 1 week.

**Validate, quantify library and sequencing**

1. Measure the concentration of the captured library using Qubit HS assay kit.
2. Measure the average fragment length of the captured library on the Tape Station.
3. Calculate molarity and multiplex for sequencing on Illumina platform.

**Consumables**:

1. xGen Hybridization and Wash Kit (Integrated DNA Technologies), 96 rxn
2. Blocking oligos - xGen Universal Blockers for Nextera libraries —NXT Mix, 96 rxn
3. Human Cot DNA (part of the Hyb and Wash Kit)
4. Ampure XP beads (Beckman Coulter Life Sciences)
5. Dynabeads M-270 Streptavidin beads (part of the Hyb and Wash Kit)
6. Eppendorf twin.tec 96 Well LoBind PCR Plates (Fisher Scientific)
7. KAPA HiFi HotStart ReadyMix (Roche)
8. Microseal B PCR Plate Sealing Film (Bio-Rad)
9. Qubit dsDNA HS Assay Kit (ThermoFisher Scientific)
10. Eppendorf tubes- 1.5 ml
11. TapeStation HS screentape and buffers (Agilent)

**Equipment:**

2. SpeedVac (ThermoFisher Scientific)
3. Magnetic Stand-96 (ThermoFisher Scientific)
4. TapeStation (Agilent)
5. Two thermal cyclers

#### Supplementary Figure S1

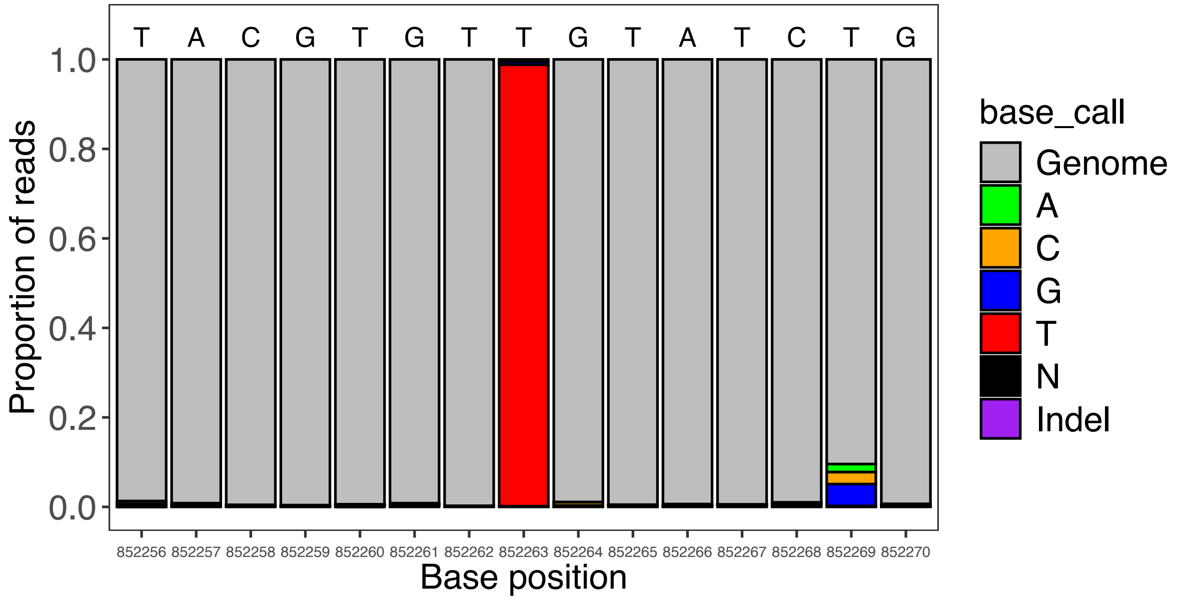

**Supplementary Figure S1.** Genomic region of Rv0757 containing the H37Ra SNP (codon 219, TCG->TTG) for one representative sample of THP-1 cells spiked with 1e6 cells H37Ra.

#### Supplementary Figure S2

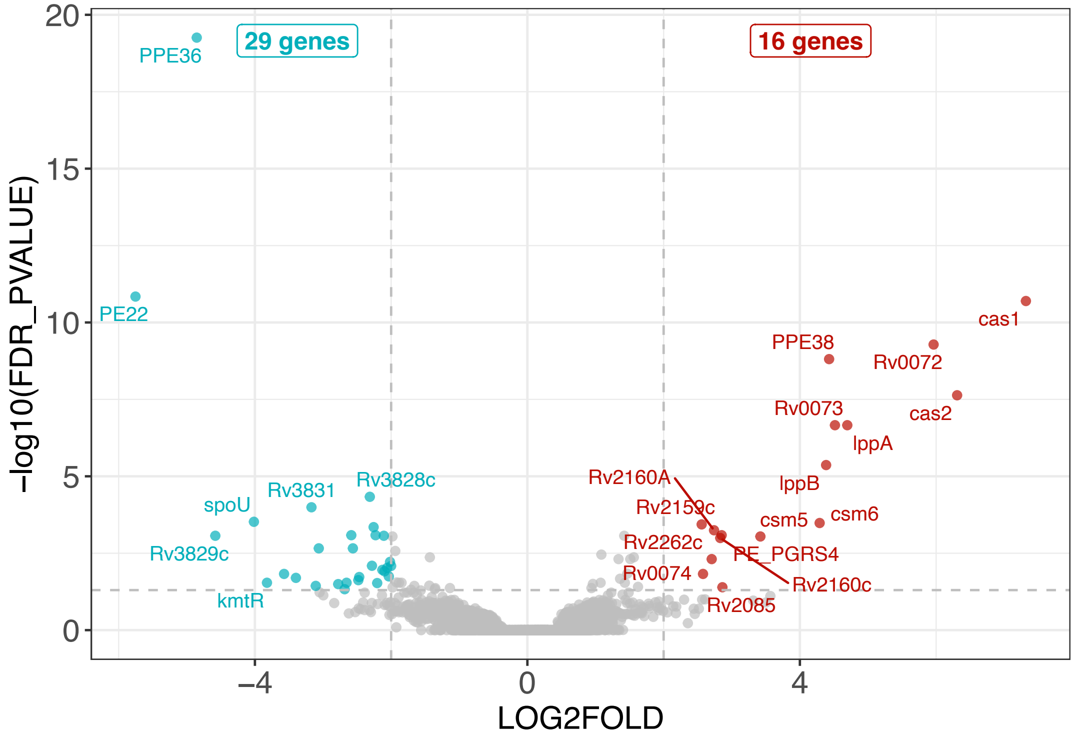

**Supplementary Figure S2.** Volcano plot of differentially expressed genes in Mtb from Lineage 4 strains (n=5) compared to Lineage 2 strains (n=4) in sputum; red points are genes significantly upregulated in Lineage 4 and blue points are genes significantly downregulated in Lineage 4 with FDR-adjusted p-value < 0.05.
